# A *Grapholita molesta* (Lepidoptera: Tortricidae) cadherin in the 99C cadherin clade binds to *Bacillus thuringiensis* Cry1A proteins

**DOI:** 10.1101/2025.11.05.686530

**Authors:** Dafne Toledo, Cristina M. Crava, Baltasar Escriche, Yolanda Bel

**Affiliations:** Laboratory of Biotechnological Control of Pests, BIOTECMED Institute, Genetics Department, University of Valencia, 46100-Burjassot, Valencia, Spain

**Keywords:** oriental fruit moth, *Cydia molesta*, Bt-cadherin, biological control

## Abstract

Bt-cadherins, a particular type of midgut cadherins, act as receptors for *Bacillus thuringiensis* Cry1A pesticidal proteins. The aim of this work was to identify and validate the Cry1A cadherin receptor in *Grapholita molesta* (GmCad1). The *GmCad1* gene was annotated by Blast genome mining using lepidopteran Bt-related cadherins. The phylogenetic analyses grouped GmCad1 with other Torticidae cadherins, in a different clade than the lepidopteran Bt-related cadherins. The *in silico* analysis of the GmCad1 showed a structure similar to that of the Bt-related cadherins, with 11 cadherin repeats (CRs), a transmembrane region, and an intracellular domain. The full-length mRNA sequencing confirmed *GmCad1* expression *in vivo* in *G. molesta* guts. To validate the binding ability of Cry1A proteins to GmCad1, a cadherin fragment (CR7-CR11) was expressed and *in vitro* binding was studied demonstrating that Cry1Aa, Cry1Ab, and Cry1Ac bind to this region in a dose-dependent manner. *In silico* molecular docking analysis suggested that the interaction may involve mainly the Domains II of Cry1Ab and Cry1Ac, and the Domain III of Cry1Aa. This study provides the first evidence of a 99-C cadherin serving as a receptor for Cry1A in *G. molesta*.

## 1. Introduction

The pesticidal Cry toxins from *Bacillus thuringiensis* (Bt) are produced and accumulated as parasporal crystals during the sporulation phase of this bacterium [1–3]. Since 3-D Cry proteins are highly specific and harmless to non-target organisms, they are used as formulated insecticides as well as expressed in transgenic plants to provide protection against insect pests, mainly caterpillars [1,2]. The most widely accepted mode of action of the Cry proteins is the pore forming model. This model has been deeply investigated mainly in Lepidoptera, leading to the following steps. The parasporal crystals are solubilized by the alkaline pH present in the caterpillar’s midgut where the proteases cleave the protein at both ends (protein activation). Subsequently, the activated proteins bind sequentially to low affinity receptors and then to midgut cadherins or/and ABC transporters with high affinity, promoting the formation of oligomeric structures which become inserted into the epithelial membrane, leading to pore formation, cell lysis, and eventually death [4–9]. How Cry proteins interact with their target receptors is closely related to their quaternary structure. Domain I (DI) is a bundle entirely formed by α-helices whose function is the pore formation in membranes. The Domain II (DII) is mainly composed of β-sheets arranged in a ‘Greek-key’ shape and involved in receptor recognition and binding. Domain III (DIII; proposed to play a role in receptor recognition, membrane insertion, and protein stability) is also formed by β-sheets in a β-sandwich conformation [1,9].

The role of cadherins as receptors for Cry proteins, particularly Cry1A toxins, has been described in various lepidopteran pest species, including *Manduca sexta* [9,10], *Heliothis virescens* [9,11], *Pectinophora gossypiella* [9,12], *Spodoptera exigua* [13], *Lymantria dispar* [9,14], *Helicoverpa armigera* [15], among others. Cadherin-like proteins have also been described in coleopteran species as receptors for Cry3 proteins, such as *Tenebrio molitor* [16] and *Alphitobius diaperinus* [17]. In addition, the Cry3 proteins are able to bind to a *Diabrotica virgifera* cadherin (DvCad1) fragment in ELISA binding assays, and DvCad1 act synergistically with the Cry3 proteins against *A. diaperinus* [18]. The importance of cadherins as receptors for Cry toxins has been demonstrated through the presence of various mutated alleles that have resulted in resistance events [11,12,20–22]. In addition, cadherin genes were knocked out using CRISPR/Cas9 technique resulting in resistance to Cry proteins in *S. exigua* [23], *H. armigera* [24], and *Ostrinia furnacalis* [25], which demonstrated the importance of cadherins in the Cry mode of action.

Cadherins constitute a large family of transmembrane glycoproteins with an important role in physiological mechanisms such as development, morphogenesis, cell recognition, signaling, and structural maintenance [9,26–28]. Structurally, all cadherins share an immunoglobulin-like β-sandwich structure and several common domains, including a transmembrane domain (TMD), an intracellular domain (ITD), and an extracellular domain. The latter varies among the different cadherin types due to the variable number of cadherin repeats (CRs) [28]. In the context of lepidopteran Bt-related cadherins, the number of CRs ranges from nine to 12 CRs, and a transmembrane ectodomain (TMED) is also present [4]. The lepidopteran Bt-related cadherins are expressed in the larval midgut epithelium [29–31] and have a molecular weight that varies depending on the species, and typically ranges from 175 to 250 kDa [4]. Previous research identified the Cry binding regions of these receptors (TBR, from Toxin Binding Region) in the extracellular region closest to the midgut membrane, which includes from CR7 to CR12 and the TMED [4,9,28].

*Grapholita molesta* (formerly *Cydia molesta*), also called the oriental fruit moth, is an invasive lepidopteran species native from Asia that belongs to the Tortricidae family. It has spread throughout temperate zones including Europe, North America, Australia, New Zealand, and South Africa [32]. *G. molesta* is an important polyphagous pest of stone and pome fruit, which has been classically controlled with chemical insecticides. However, this species has developed resistance to ten organophosphates and three carbamates, reducing the number of active ingredients available for its management [33]. As an alternative, pheromones [33], pheromone synergistic chemicals [33], entomopathogenic fungi [34] or parasitoids [35] have been used in an attempt to control this pest. Few studies have been published about the effects of Bt Cry proteins on *G. molesta*, but it has been reported to be highly susceptible to both activated Cry1Aa and Cry1Ac toxins, representing promising results for the use of this technique in the field control of *G. molesta* [36]. However, there is still a gap of knowledge regarding the crucial events of the mode of action of Cry1Aa and Cry1Ac in *G. molesta*. For example, it is unknown which proteins act as receptors for these two toxins.

In this study, we present the first description of a *G. molesta* cadherin that acts as a Cry1A receptor *in vitro* (GmCad1). Its presence was predicted *in silico* and later confirmed in *G. molesta* guts. The binding ability of Cry1Aa, Cry1Ab and Cry1Ac Bt proteins to GmCad1 was verified using a cadherin fragment comprising CR7-CR11. The residues that could be involved in the interactions were predicted using a molecular docking approach. The phylogenetic relationship of the protein was analyzed using lepidopteran and coleopteran cadherins. The results shown in this work evidenced that the GmCad1 could act as a receptor for Cry1A toxins in *G. molesta*.

## 2. Materials and Methods

### 2.1. Cadherin gene mining

The *G. molesta* cadherin gene sequence was identified through mining the genome of *G. molesta* [37], which was assembled at the chromosome level (BioProject ID: PRJNA627114). Iterative TBLASTN [38] searches were conducted using the amino acid sequence of Bt-related cadherins from Ostrinia nubilalis (Acc. Number: AAY44392.1), S. exigua (Acc. Number: AGS80251.1), and Bombyx mori (Acc. Number: BAA77212.1) as queries. The best hit obtained from these TBLASTN searches allowed the identification of the *G. molesta* chromosome containing the cadherin sequence. To get this sequence, the whole chromosome was annotated using the FGENSH tool [39] (http://www.softberry.com/), which predicts exon/intron junctions, mRNA, and protein sequences. The predicted proteins obtained from FGENSH analyses were filtered to include lengths between 1200 and 1900 amino acids. These filtered proteins were later used as queries in BLASTP searches [38] against the whole nr database, to identify sequences with homology to Bt-related cadherins (specifically containing from 9 to 12 CRs).

### 2.2. Synthesis of cDNA

*G. molesta* last instar larvae, obtained from a colony reared at the University of Valencia [36], were dissected and their guts isolated. The guts were washed in Phosphate-Buffered Saline (PBS; Fisher bioreagents, Geel, Belgium), and placed in tubes containing 50 µl of RNALater® (Merck, Darmstadt, Germany).

Total RNA was purified from dissected guts using the RNeasy Minikit (Qiagen GmbH, Hilden, Germany) following manufacturer’s instructions. The RNA samples were treated with DNase I (Thermo Fisher Scientific Baltics UAB, Vilnius, Lithuania). Subsequently, the RNA was retrotranscribed into cDNA using PrimeScriptTM RT Reagent Kit (Takara Bio Inc. Kusatsu, Japan) by RT-PCR in a thermal cycler (Eppendorf Mastercycler; Eppendorf, Hamburg, Germany) according to the manufacturer’s instructions. The obtained cDNA was aliquoted, frozen in liquid nitrogen, and stored at-80 °C until used for the amplification of the GmCad1 sequence.

### 2.3. GmCad1 amplification and sequencing

Several primer pairs were designed to amplify the entire coding sequence of GmCad1. PCR amplifications were conducted using KAPA HiFi DNA polymerase (KAPA Biosystems Pty, Cape Town, South Africa) following the manufacturer’s instructions, with 100 ng of cDNA as template. The PCR products were visualized after electrophoresis on 1 % agarose gels. Once the size of the amplicon matched the expected size, the band in the gel was excised, purified using NucleoSpin® Gel and PCR Clean-up kit (Macgeret-Nagel GmbH & Co., Düren, Germany) and sequenced using the Sanger method at Stab Vida facilities (Investigação e Serviços em Ciências Biologicas, LDA, Portugal). The obtained sequences were aligned to the predicted nucleotide sequence using the Unipro UGENE program [40]. A final consensus sequence was generated using at least two sequenced fragments at any given position to ensure accuracy. The sequence was deposited at NCBI with Acc. Number OQ943817.

### 2.4. In silico study of the GmCad1 sequence

The real exon/intron junctions of the GmCad1 gene were obtained by aligning the whole coding sequence obtained using Sanger sequencing with the *G. molesta* chromosome sequence (BioProject ID: PRJNA627114) using Unipro UGENE software [40]. The GmCad1 coding sequence was translated to protein sequence with the same program and analyzed with the Simple Modular Architecture Research Tool [41,42] (SMART, http://smart.embl-heidelberg.de/) to search for the conserved domains/features.

The phylogenetic relationship of GmCad1 with other lepidopteran cadherins was assessed aligning their amino acid sequences with the Clustal Omega tool (https://www.ebi.ac.uk/Tools/msa/clustalo/) from the European Bioinformatics Institute facilities. The phylogenetic tree was built employing the MEGA11 software [43] using the Maximum Likelihood method and Poisson correction model with 150 bootstraps. The resulting phylogenetic tree was modified with iTOL [44] (https://itol.embl.de/) for visualization.

### 2.5. Cloning, expression, and purification of CR7-CR11 fragment

The nucleotide sequence corresponding to the CR7-CR11 segment of the GmCad1 was selected based on the GmCad1 protein quaternary structure obtained from SMART program. The corresponding nucleotide sequence was optimized for Escherichia coli expression, synthesized by GeneScript (Leiden, Netherlands) including an N-t Histidine-tag and kanamycin (kan) resistance gene, and cloned into the pET-30a(+) vector. The plasmids were transformed into BL21 (D3) E. coli competent cells by heat-shock transformation [45]. Briefly, 0.4 µg of the plasmid were added to the competent cells. After 20 min on ice, the cells were incubated for 90 s at 42 °C and left on ice for 5 min. To recover the cells, 900 µl of Luria Bertani broth (LB; 10 g/l NaCl, 10 g/l tryptone, 5 g/l yeast extract) was added and incubated for 1 h at 37 °C, with shaking at 180 rpm. The cell suspension was centrifuged at 6000 xg for 1 min at room temperature. The resuspended pellet was plated on LB-agar supplemented with 50 µg/ml kanamycin and incubated for 16 h at 37 °C. A negative heat-shock control was performed by using the same procedure but adding 2 µl of sterilized Milli-Q water to the E. coli competent cells.

The CR7-CR11 expression and purification were performed as described by Ren et al. [31] with some modifications. One transformed colony was inoculated into 7 ml of LB medium supplemented with 50 µg/ml kanamycin and incubated for 16 h at 37 °C, 180 rpm. From this preculture, 1 ml was inoculated into 700 ml of LB supplemented with 50 µg/ml kanamycin. The culture was incubated at 37 °C, 180 rpm, until the optical density value (λ= 600 nm, OD600) reached 0.5-0.8. The protein expression was then induced with 0.8 mM IPTG (isopropylthiol-β-D-galactoside; Fisher bioreagents, Geel, Belgium), and the culture was incubated for 16 h at 25 °C, 180 rpm. The cells were pelleted by centrifugation for 20 min at 4 °C, 15000 xg and kept for 4.5 h at-80 °C. The pellet was thawed and resuspended in 20 mM Tris-HCl, 7.4 pH, sonicated, and centrifuged for 15 min at 4 °C, 17420 xg. The pellet was resuspended in 20 mM Tris-HCl, 2 M urea, 0.5 M NaCl, 2 % Triton X-100, 7.4 pH, sonicated and centrifuged as above. To solubilize the protein of the E. coli inclusion bodies, 20 ml of 20 mM Tris-HCl, 6 M Urea, 0.5 M NaCl, 5 mM Imidazole, 0.5 mM β-mercaptoethanol, 7.4 pH were added to the pellet, and the suspension was incubated for 1 h with gentle shaking. After centrifugation under the same above conditions, the supernatant was loaded onto a HisTrap TM FF crude column (GE Healthcare Bio Sciences, Uppsala, Sweden). A decreasing urea gradient was used to refold the peptide bound to the column matrix to its native structure. The rest of purification steps were performed according to the manufacturer’s instructions. The purified peptide fractions were checked by electrophoresis on 12 % SDS-PAGE gels and quantified by the Bradford method [46] using BSA (Bovine Serum Albumin) as standard.

### 2.6. Preparation of Cry1A proteins and dot blot binding assays

The Cry1Aa and Cry1Ab proteins were obtained from recombinant E. coli strains kindly provided by Dr. R. de Maagd (Wageningen University, Nederlands). The proteins were produced, solubilized, and activated as described elsewhere [47]. The Cry1Ac was prepared from a recombinant Bt strain (Ecogen In., Langhorn, PA, USA) according to the published methodology [48].

The dot blot binding assays were performed following Fabrick and Tabashnik [49] with some modifications. Increasing amounts of CR7-CR11 fragment (0.125 µg, 0.25 µg, 0.5 µg, and 1 µg) were dotted onto AmershamTM ProtanTM 0.45 µm NC 200 mm x 4 m Nitrocellulose Membrane (Fisher Scientific, Madrid, Spain), followed by membrane blocking with PBS-B (PBS, 5% skimmed milk) for 16 h at 4 °C. The membrane was then incubated with 0.1 µg/ml of Cry1A trypsin-activated protein (Cry1Aa, Cry1Ab or Cry1Ac) in PBST (PBS, 0.1% Tween-20) for 1 h, followed by three washes with PBST (ten minutes each). The membrane was incubated with the polyclonal α-Bt Cry1A antibody (Eurofins Abraxis, Warminster, England) at a 1:5000 ratio in PBST 0.5 % skimmed milk for 1 h, and washed with PBST as before. The α-rabbit IgG-conjugated horseradish peroxidase antibody (Sigma-Aldrich, St. Louis, USA), at a 1:10000 ratio, was added to the same solution and incubated for 1 h. After three washes with PBST, the membranes were revealed with SuperSignal® West Femto Maximum Sensitivity Substrate (Thermo Scientific, Rockford, IL, USA) according to the manufacturer’s protocol, and visualized with ImageQuant LAS400 (GE Healthcare BioSciences, Uppsala, Sweden). As positive controls, 0.1 µg of each one of the Cry1A proteins were blotted on the membrane. The negative control consisted in blotting the same amounts of CR7-CR11 peptide used in the experiments into the membranes and incubating them with just PBST. Three replicates were carried out for each assay. The images were contrast adjusted using the PhotoScapeX program, with all pixels equally adjusted from a treshold of 50 to 65.

### 2.7. Modelling of the CR7-CR11 protein fragment structure

The GmCad1 amino acid sequence was uploaded to the Phyre2 server [50] (http://www.sbg.bio.ic.ac.uk/) to obtain its 3-d structure. The sequence from the CR7 to the CR11 (see section 2.4) was selected using the Chimera 1.16 software [51] for the subsequent bioinformatic analyses. The quality of the CR7-CR11 predicted structure was evaluated by a Ramachandran’s plot generated with PROCHECK [52,53] in the SAVES v6.0 server (https://saves.mbi.ucla.edu/). The resulting predicted structure was refined employing the ModRefiner tool [54] (https://zhanggroup.org/ModRefiner/) and compared to the initial model by the values of its Ramachandran’s plot.

### 2.8. Molecular docking and interaction prediction

The molecular docking was performed with the refined structure of the CR7-CR11 fragment. Regarding the Cry1A proteins, the Cry1Aa X-Ray diffraction protein structure (PDB ID: 1CIY) [55] was used and the PDB structures of Cry1Ab and Cry1Ac were obtained from AlphaFold [56,57]. The quality of the Cry1Ab and Cry1Ac models was analyzed through Ramachandran’s plot using the same tool as described above. The Cry1Aa, Cry1Ab, Cry1Ac, and GmCad1 protein structures were submitted to the H++ server [58–60] (http://newbiophysics.cs.vt.edu/H++/) to obtain the structures with the missing hydrogens and their amino acid protonation states at different pH values (5, 7.5, and 10.5). The salinity, internal dielectric, and external dielectric values were set as default. The prediction of the binding conformation between the CR7-CR11 and Cry1A toxins was performed on the HawkDock server [61–63] (http://cadd.zju.edu.cn/hawkdock/). The models were re-ranked by the MM/GBSA utility [61,64–66] on the same webpage. The top-ranked model for each toxin-receptor interaction was selected for further analysis. The LigPlot+ v2.2 program [67,68] was used to plot the Cry1A-receptor interactions.

## 3. Results

### 3.1. G. molesta cadherin gene mining and protein sequence prediction

The TBLASTN searches revealed that chromosome 4 (Acc. Number CP053123.1) of the *G. molesta* genome likely harbored genes encoding cadherins, characterized by having nine to 12 CR repeats. Subsequent gene annotation, length filtering of *in silico* predicted sequences, and additional BLASTP searches identified two hypothetical proteins on chromosome 4 with features compatibles with Bt-related cadherins. One of the identified cadherins has a length of 1744 amino acids and ten CRs. It shows high homology with the 89-D cadherin clade in a BLASTP search. The other cadherin had 1612 amino acids and 11 CRs, and showed high homology with the 99-C cadherin clade, as will be discussed later. We chose the latest one because several Bt-related cadherins in lepidopteran species have the same number of CRs, and for subsequent analysis we focused on this cadherin. We named GmCad1, and it was encoded by the locus between nt 11156887 and nt 11176471. *In silico* prediction of the GmCad1 CDS yielded a length of 4839 nt, encoding a hypothetical protein of 1612 amino acids. Intron/exon detection showed that the gene comprised 29 exons (Table 1). The GmCad1 protein sequence exhibited the highest homology in BLASTP searches against the whole nr database with a 99-C cadherin predicted by automated computational analysis from the genome of *Leguminivora glycinivorella* (Acc. Number: XP_047998198.1), a lepidopteran species belonging to the same family as *G. molesta*, with 91% identity, 95% positive, and 0% gaps.

**Table 1.**
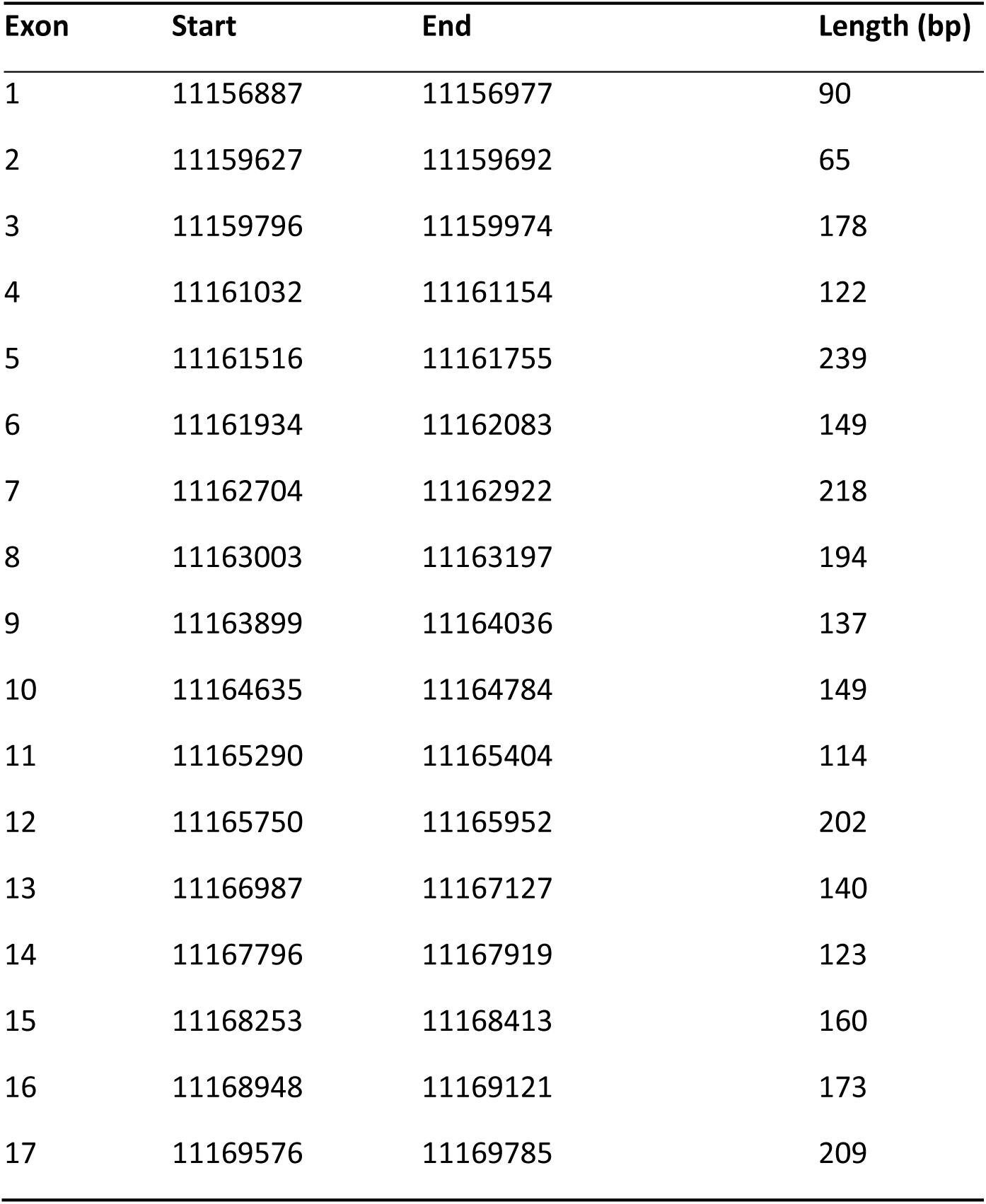
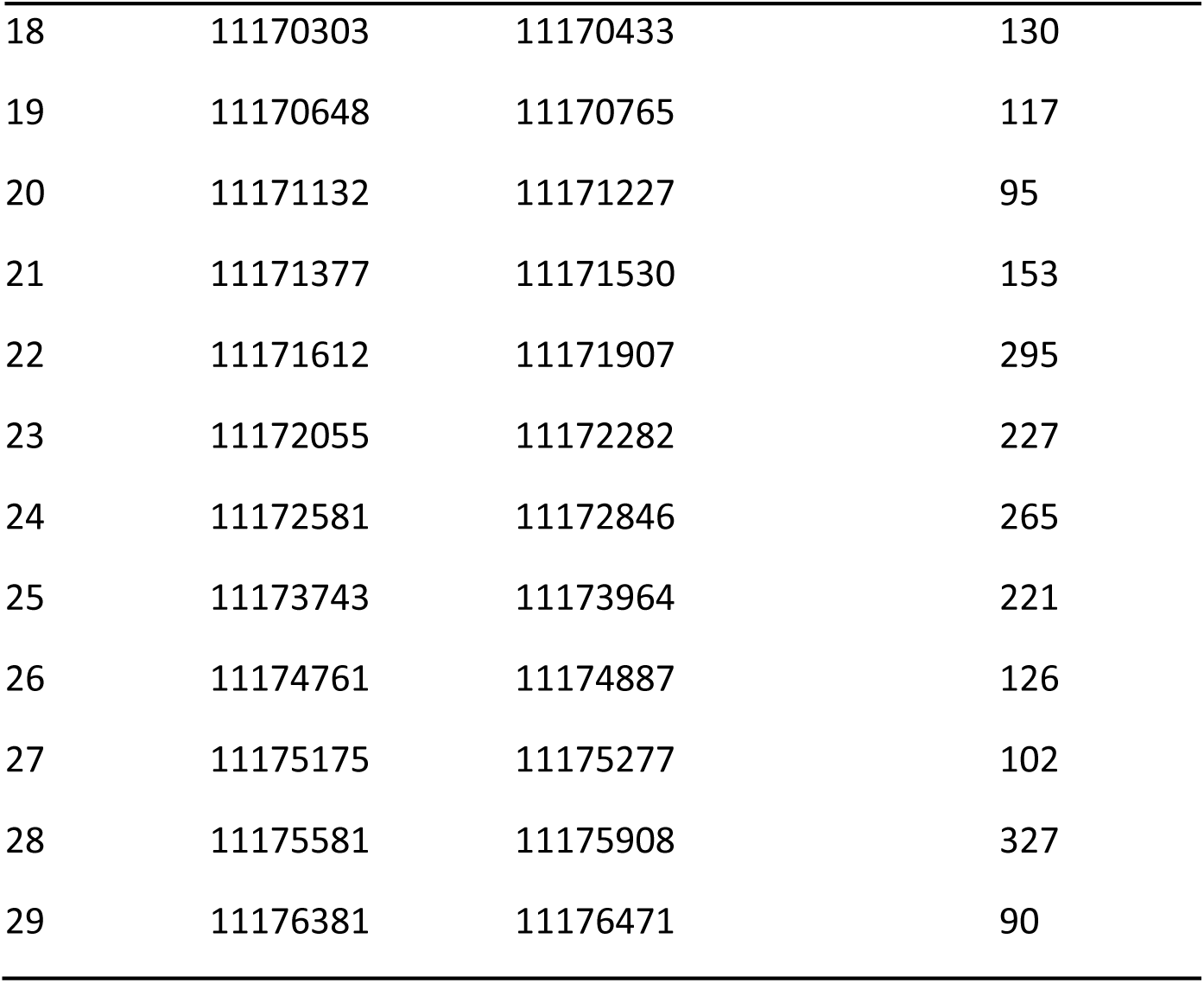
*G. molesta* cadherin gene structure. The table shows the coding regions (exons) and its location within the fourth chromosome of the *G. molesta* genome (BioProject ID: PRJNA627114).

### 3.2. G. molesta cadherin transcript sequencing and in silico study of the predicted protein

The entire CDS of the *GmCad1* gene was amplified from cDNA obtained from last instar larval guts, confirming its expression in this compartment. The experimentally obtained sequence matched the predicted length from the whole *G. molesta* genome analysis. The alignment of the sequenced CDS with the predicted one (see section 3.1) revealed 53 single nucleotide polymorphisms. Of these, only five were non-synonymous, resulting in amino acid changes (Table 2). *In silico* analysis of the translated protein revealed that GmCad1 contained conserved domains typical of Bt-related cadherins: a small SP, 11 CR, a TMED region, a TM domain, and a low complexity sequence that may correspond to the protein’s ITD (Figure 1). The phylogenetic analysis of GmCad1 involved several Bt-related cadherins from Lepidoptera and Coleoptera, along with another type of insect cadherins known as the 99-C cadherins. These 99-C cadherins share structural similarities with Bt-related cadherins but have not been implicated in Bt mode of action. The results (refer to Figure 2) revealed that GmCad1 grouped within the 99-C cadherin clade rather than with the lepidopteran Bt-related cadherin clade, supporting that GmCad1 is not orthologous to other cadherins involved in Bt-mode of action. Specifically, GmCad1 clustered with cadherins from other species within the Tortricidae family (such as *L. glycinivorella* and *Choristoneura fumiferana*), as well as with other lepidopteran 99-C cadherins. Notably, the discovery of the GmCad1 sequence through iterative BLAST searches using Bt-related cadherins, coupled with its clustering pattern, suggests that cadherins similar to the Bt-related ones described in lepidopterans could be absent in *G. molesta*.

**Figure 1.**
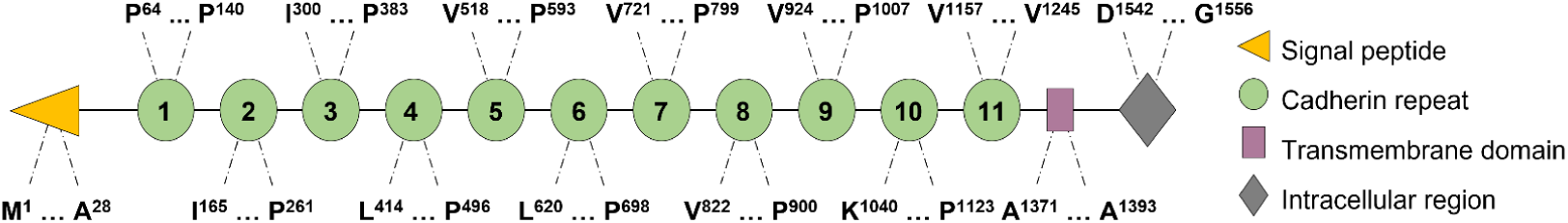
Schematic representation of the *G. molesta* cadherin protein. The length of each conserved domain is indicated by the amino acid positions from left to right.

**Figure 2.**
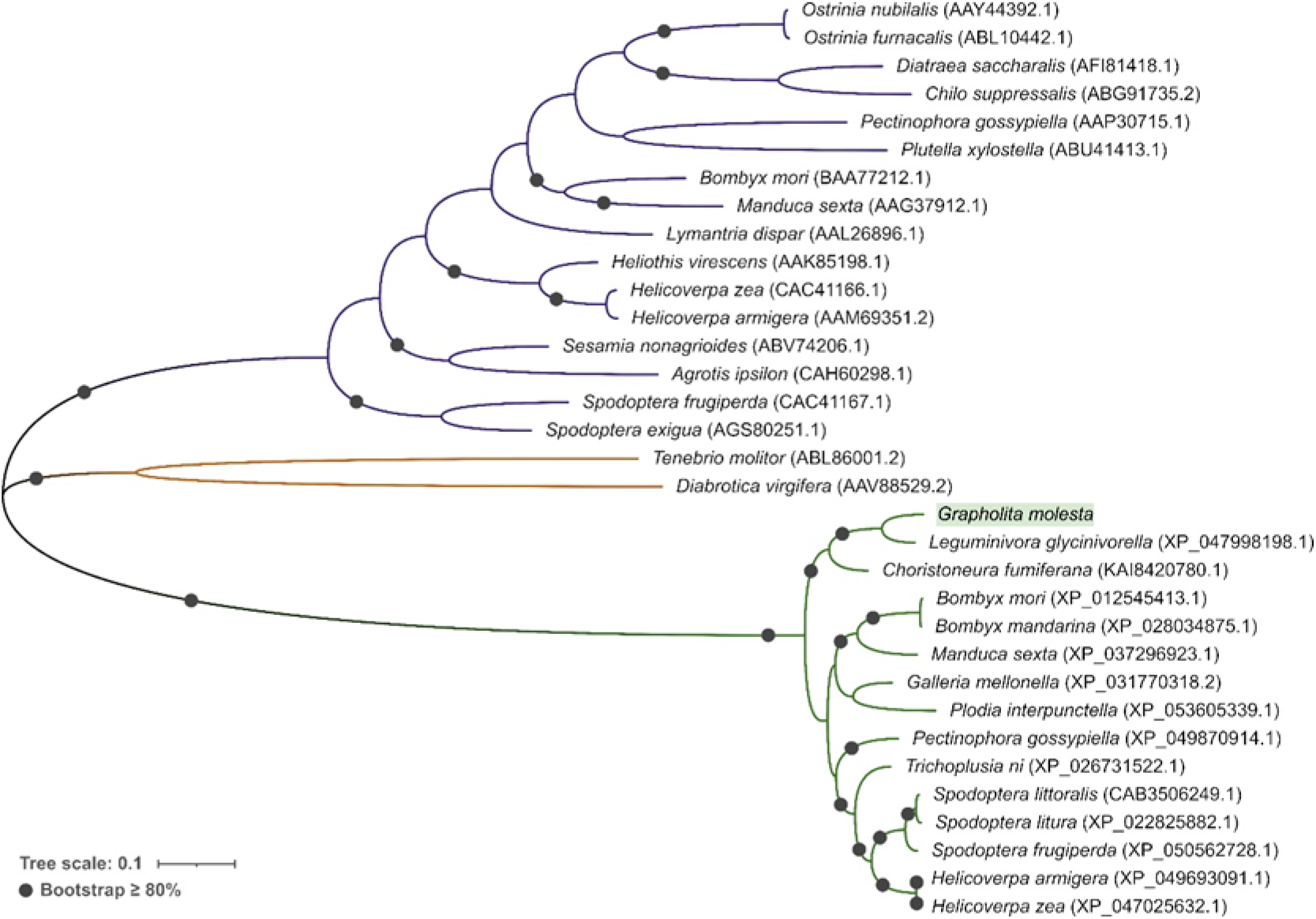
Unrooted phylogenetic tree of insect cadherin proteins, inferred using the Maximum Likelihood method and Poisson correction model with 150 bootstraps. Evolutionary analysis was conducted in MEGA11 and modify with iTOL. The NCBI accession numbers of the proteins are shown in brackets. The described lepidopteran Bt-related sequences are shown by violet lines, the coleopteran ones by brown lines, and lepidopteran 99-C cadherins similar ones by green lines.

**Table 2.**
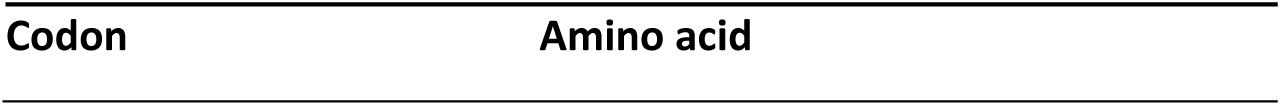
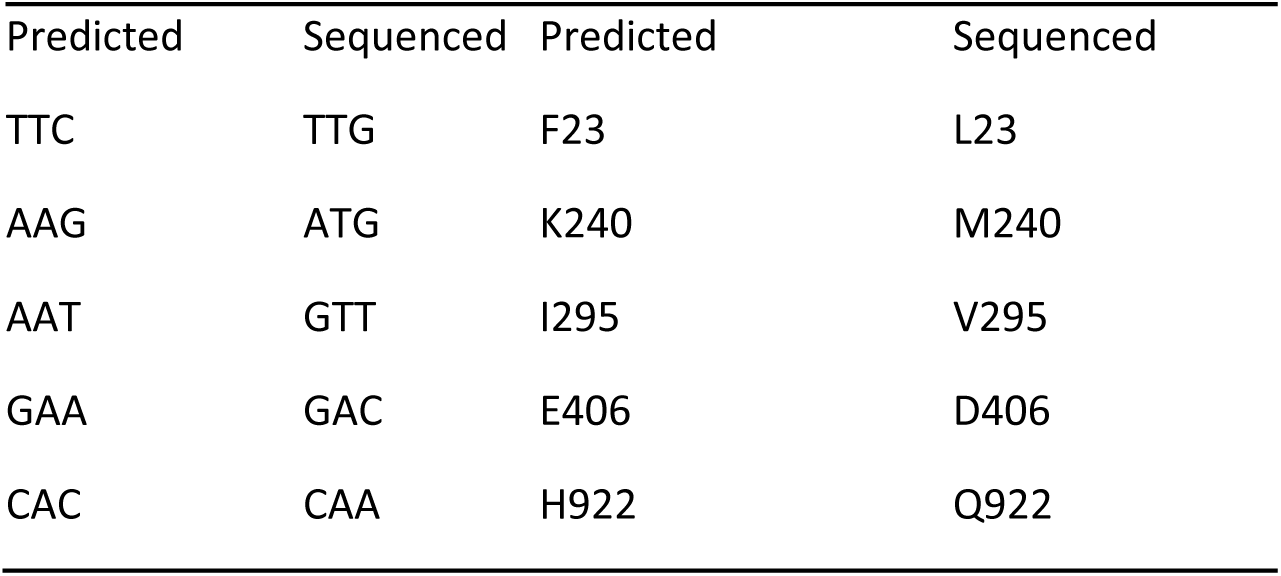
Codon changes between the predicted sequence based on the released genome (BioProject ID: PRJNA627114) analysis and the experimentally sequenced *GmCad1* (Acc. Number: OQ943817) gene that involve an expected change in the amino acid sequence.

| Predicted | Sequenced | Predicted | Sequenced |
| --- | --- | --- | --- |
| TTC | TTG | F23 | L23 |
| AAG | ATG | K240 | M240 |
| AAT | GTT | I295 | V295 |
| GAA | GAC | E406 | D406 |
| CAC | CAA | H922 | Q922 |

### 3.3. Binding of Cry1A proteins to GmCad1

In order to validate experimentally the ability of GmCad1 to bind to Cry1A protein, we expressed and purified the CR7-CR11 GmCad1 fragment, which should harbor the toxin biding region (TBR). The SDS-PAGE electrophoretic analysis showed a good expression, as a band of the expected molecular weight of ∼70 kDa (Figure 3a).

**Figure 3.**
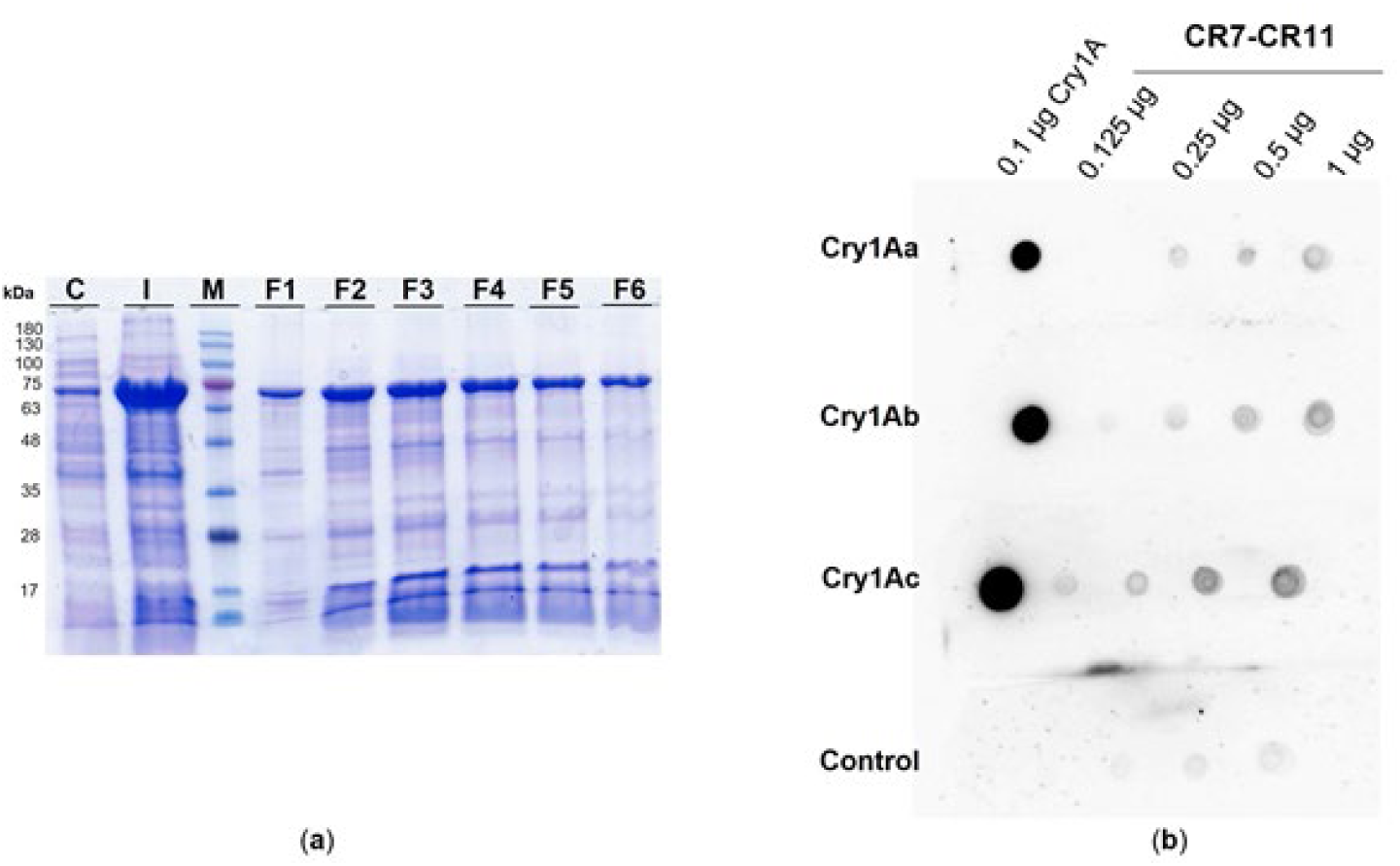
Expression, purification and Cry1A binding analyses of the GmCad1 CR7-CR11 fragment. (**a**) SDS-PAGE analysis of the fractions obtained after column purification of the peptide expressed in *E. coli* cells. (**b**) Cry1A dot blot binding assay of the Cry1 proteins to the CR7-CR11 peptide. C: 10 µl of the Gmcad1 CR7-CR11 culture. I: 10 µl of the column input. M: Blue Star Marker (Nippon Genetics Europe GmbH, Düren, Germany). Lanes labeled from F1 to F6 are 10 µl of the eluted fractions. Control: GmCad CR7-CR11 without incubation with Cry1A proteins.

Dot blot binding assays were performed to test the ability of Cry1Aa, Cry1Ab and Cry1Ac proteins to bind to the refolded CR7-CR11 peptide in non-denaturing conditions. The results showed binding in a dose-dependent manner for the three Cry proteins. A slight signal was observed in the negative controls due to a cross-recognition of the antibody used, but the different signal intensities of the assays regarding the negative control evidenced specific binding of each Cry1A protein to the CR7-CR11 cadherin fragment (Figure 3b).

### 3.4. Toxin-receptor molecular docking

The interaction between Cry1A and GmCad1 was analyzed *in silico* through molecular docking. For this analysis the predicted structure of the *G. molesta* cadherin was obtained by homology with six different protocadherin structures (PDB IDs: 6E6B, 6VG1, 6VG4, 6C13, 5SZN, 5T9T). The structure of CR7-CR11 was submitted to ModRefiner to improve the quality of the model. The Ramachandran’s plot of the initial structure (Figure 4a) showed 88.1% of the residues in the most favorable regions and 9.6% in additional allowed regions. These values were increased in the refinement (Figure 4b), obtaining 90.6% of the residues in the most favorable regions and 8.7% in additional allowed regions. Since the refined structure contained over 90% of residues in the most favorable positions, the model can be considered as good for further molecular docking analysis.

**Figure 4.**
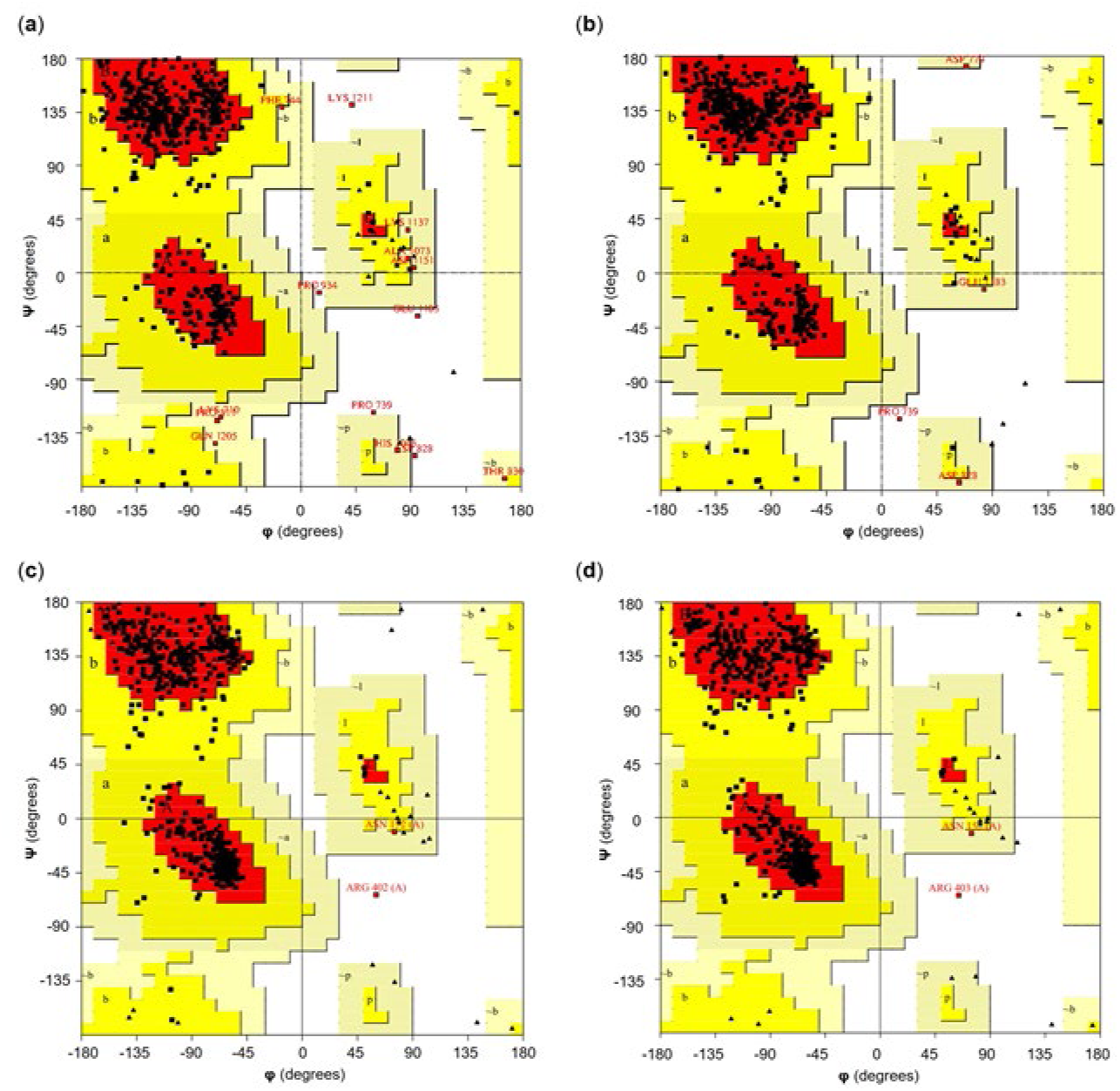
Ramachandran’s protein plots obtained by PROCHECK analysis. (**a**) CR7-CR11 initial model. (**b**) CR7-CR11 refined model. (**c**) Cry1Ab. (**d**) Cry1Ac.

For all three Cry1A toxins, the results obtained for the molecular docking and its binding free energy, as well as the amino acids that are interacting with the GmCad1 at pH acidic (pH 5), neutral (pH 7.5) or basic (pH 10.5) were exactly the same. For this reason, we evaluated the results at pH 10.5 because this is the condition that most resembles that of the lepidopteran midgut where the interaction between Cry1A proteins and GmCad1 receptor occurs.

The molecular docking approximation of the Cry1Aa protein and the receptor (Figure 5a) had a-56.38 kcal/mol binding free energy. This in silico value indicated that the binding between CR7-CR11 and Cry1Aa toxin was theoretically strong. The LigPlot+ graph showed that the Cry1Aa protein interacts mostly with the CR10 (from amino acid K1040 to P1123), and the residues that may be important in the interaction were mostly located in the β21-β22 junction (N578 and S580) and β16 (R511) of the protein’s DIII. Two residues from DII, S351 and T353 were also found that could interact with the CR10 of the GmCad1.

**Figure 5.**
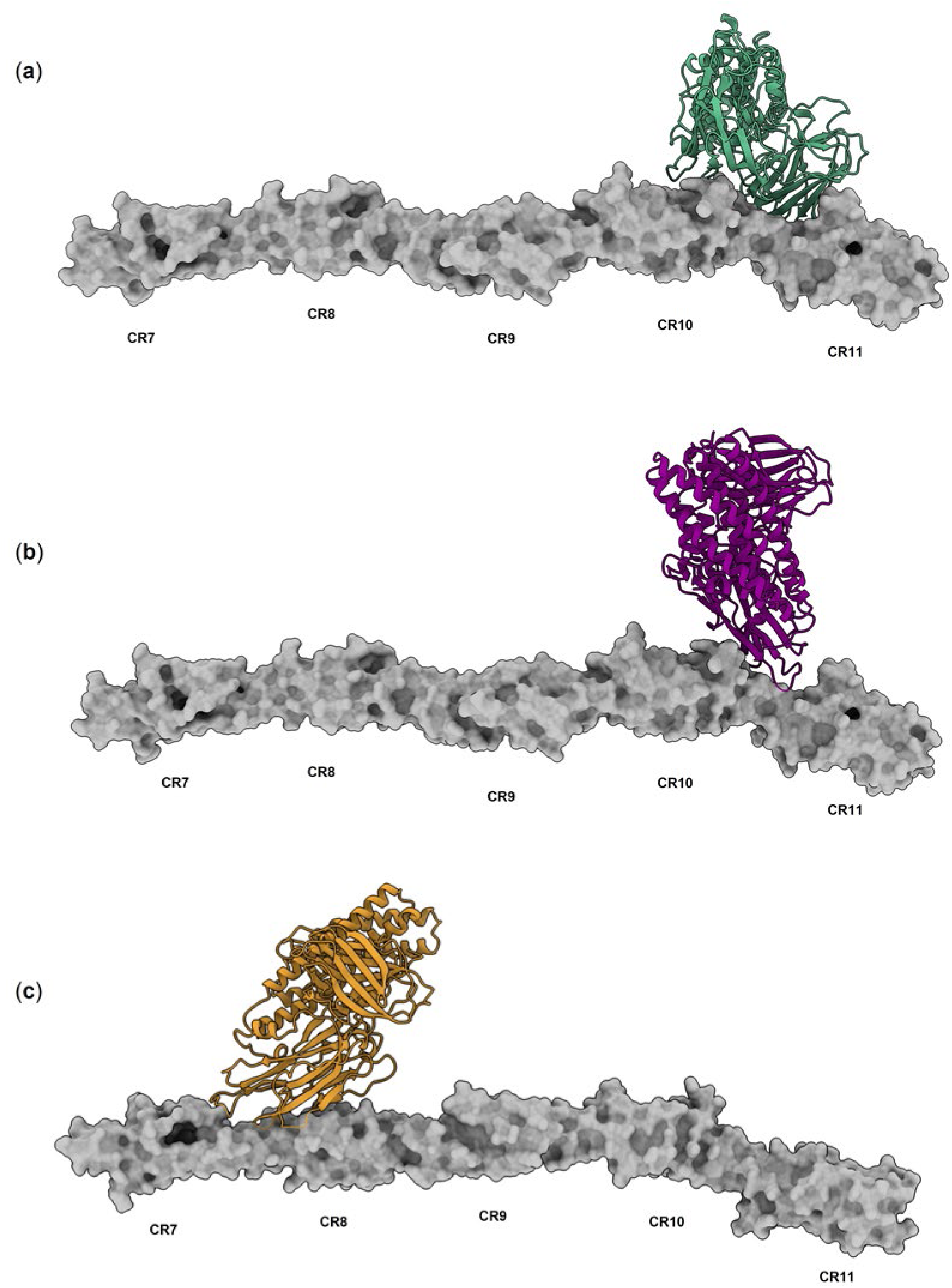
Molecular docking results of the Cry1A proteins and the CR7-CR11 from *G. molesta* at pH 10.5. (**a**) Cry1Aa. (**b**) Cry1Ab. (**c**) Cry1Ac. The image of the complex was obtained with the Chimera X software.

The structure quality of the Cry1Ab protein was analyzed by Ramachandran’s plot (Figure 4c), which displayed that more than 92% of all residues were in the most favorable regions. The Cry1Ab and CR7-CR11 interaction complex (Figure 5b) had a-57.38 kcal/mol binding free energy. The interaction diagram corroborated that mostly loop 1 (R311) and loop 2 (R368 and N372) amino acids, as well as E313 (β3r sheet of DII) could interact with several residues of the CR10 and with amino acids present in the junction between the CR10 and CR11 (from amino acid P1123 to V1157).

Regarding to Cry1Ac structure, 92.4% of the residues were located in the most favored conformations (Figure 4d). The Cry1Ac-receptor complex (Figure 5c) had a-58.23 kcal/mol of binding free energy. The interaction diagram provided by LigPlot+ showed that the Cry1Ac protein mostly interacted through amino acid R311 of the loop 1 and β8-β9 junction amino acids R403, Q404 and R405 of the DII with the cadherin domains from CR7 to CR8 (from amino acid V721 to P900).

## 4. Discussion

Since its discovery in 1901, Bt has been extensively investigated for its potential as a bioinsecticide, leading to the development of formulations and transgenic plants expressing Bt proteins [1,2]. Despite over a century of intensive research, the mode of action of the Bt 3-D Cry proteins remains incompletely understood [69,70]. It is generally assumed that cadherin receptors promote the formation of the Cry protein pre-pore complex [71], a critical step in their toxicity, highlighting the importance of this family of midgut receptors. Studies conducted thus far indicate that cadherin receptors may contribute differently to toxicity depending on the specific Bt toxin and the target pest [13,31,72–74]. Therefore, characterizing this group of receptors in lepidopterans, the main targets of Cry toxins, is key for understanding the specific mode of action of a toxin in a given insect species. In this study, we aim to provide further insight into the role of cadherin receptors in the mode of action of Cry1A proteins in the lepidopteran pest *G. molesta*, which is susceptible to Bt Cry1Aa and Cry1Ac proteins. We describe, for the first time, a cadherin protein belonging to the 99-C group as a candidate receptor for the toxic Bt proteins in this insect. The *G. molesta* cadherin gene was identified by TBLASTN searches using the *O. nubilalis*, *S. exigua*, and *B. mori* Bt-related cadherins amino acid sequences as a query. The predicted cadherin gene, located on the chromosome four of *G. molesta*, has a coding region of 4839 bp length and encodes a protein of 1612 amino acids.

Bt-related cadherins are mostly expressed in the midgut epithelium of insects [29–31]. The cDNA used for the gene sequencing in the present work was obtained from whole guts, confirming the presence of the *G. molesta* cadherin transcripts in the tissue targeted by Bt Cry proteins. After characterizing the *G. molesta* cadherin gene and protein, the ability of the Cry1A proteins to bind to this putative receptor was tested by means of dot blot binding assays. These assays showed that Cry1Aa, Cry1Ab and Cry1Ac were able to bind to the refolded CR7-CR11 GmCad1 fragment in non-denaturing conditions in a doses-dependent manner.

Although the number of CRs of GmCad1 is consistent with that of the cadherins described so far as receptors for the Bt toxins in several lepidopteran insects such as *P. gossypiella*, *H. virescens*, *O. nubilalis* and *S. exigua* [4], and our *in vitro* experiments demonstrated its ability to bind Cry1A proteins, phylogenetic analysis revealed that it clustered with 99-C cadherins. These 99-C cadherins possess divergent amino acid sequences from Bt-related cadherins, although they maintain high structural similarity [75]. Thus far, 99-C cadherins have not yet implicated in the mode of action of Bt in any lepidopteran or coleopteran species. Notably, a recent study identified another divergent cadherin gene involved in the mode of action of the Cry proteins in *H. zea*, belonging to the 86-C clade. In this species, mutated alleles of the 86-C cadherin gene conferred up to 10-fold resistant to Cry1Ac protein. Despite this association was established, the mechanisms of resistance in this species were not completely explained by the changes in the 86-C cadherin [76]. Together with our results, this suggests that the range of cadherins involved in the mode of action of Bt may be broader than previously thought, with other cadherin types outside the Bt-related clade potentially acting as receptors, indicating possible redundant mechanisms within certain subfamilies of cadherins. In fact, the silencing of the *Chilo suppressalis* cadherin CsCad1 (which grouped with the Bt-related cadherins), or the silencing of CsCad2, CsCad3, and CsCad4 (which grouped in other cadherin clades) decreased the susceptibility to Cry1 or Cry2 proteins [77,78] showing that different genes of the cadherin family are involved in the mode of action of Cry toxins in this lepidopteran species. In contrast, in *Plutella xylostella*, only silencing of the Bt-related cadherin PxCad1 conferred resistance to Cry proteins, unlike silencing of five other cadherin genes [79]. These studies suggest that diverse Cry cadherin receptors may have different importance in the Cry mode of action depending on the species. This scenario resembles that of the ABC transporters, a large family of membrane receptors implicated in some Cry toxin resistance events, which appear to have a redundant role as Cry receptors [8]. Indeed, studies have shown that several Cry proteins (including Cry1Aa, Cry8Ca, Cry1Da, Cry9Aa, and Cry1Ca) can bind to multiple ABC receptors in *B. mori*, demonstrating a promiscuous binding to this family of receptors [80]. These findings underscore the challenges in determining the role of the receptors in the mode of action of several Bt proteins, their potential redundant roles, and the large number of possible candidates within the same receptor family.

A computational molecular docking analysis was performed to predict how the Cry1A proteins could interact with the putative toxin binding region of GmCad1, which in Bt-related cadherins involves CR7-CR11. This approach has been previously used to characterize the binding regions between Cry1Fa [81] and Cry1Ac [82] to *P. xylostella* cadherin. Following the molecular docking simulation, the protein-receptor complexes were re-ranked and analyzed, obtaining consistent results for the interactions at pH levels of 5, 7.5, and 10.5. The results showed that the Cry1Aa protein interacted with the CR10-TMED region of GmCad1 by its DII and DIII domains. Similarly, Cry1Ab interacted with almost the same region of the GmCad1, predominantly through its DII loop regions. In contrast, the molecular docking results for Cry1Ac indicated potential interaction with the CR7-CR8, mediated by its DII loop 1 and the flexible region between the β8 and β9 of the same domain. The varying binding positions predicted for each Cry1A protein to GmCad1 were not unexpected, given previous evidence of Cry1A binding to several lepidopteran Bt-related cadherin regions [7,15,49]. While the most commonly described TBRs are typically located in CR7, CR11, CR12 and TMED [4,49,83–86], interactions with other non-canonical TBRs have also been reported. For instance, Cry1Ac toxin has been shown to interact with the CR8-TMED of the *P. gossypiella* cadherin [49], and Cry1Fa was predicted by molecular docking to bind the CR10 region of *P. xylostella* cadherin, later confirmed by Elisa binding assays [81]. Additionally, a deletion of 36 amino acids in the CR5 of *P. gossypiella* cadherin was linked to Cry1Ac resistance [87].

In summary, the present work evidences the role of the *G. molesta* cadherin GmCad1 as possible putative receptor for the Cry1A proteins. This study contributes to the knowledge of the mode of action of cry proteins and opens the possibility of an alternative cadherin receptor for these proteins. Future studies will focus on the role of the 99-C cadherin in other insect species, and will help to design the best strategies for the use of Bt proteins to control lepidopteran pests and to manage the resistance to Cry proteins.

## Author Contributions

Conceived and designed the experiments: DT, YB, and BE. Performed the experiments: DT, YB, and MCC. Analyzed the data: DT and YB. Wrote the original draft: DT and YB. Wrote - reviewed & edited: DT, YB, and BE. Finding and project administration: BE.

## Supplementary Materials

**Figure S1.**
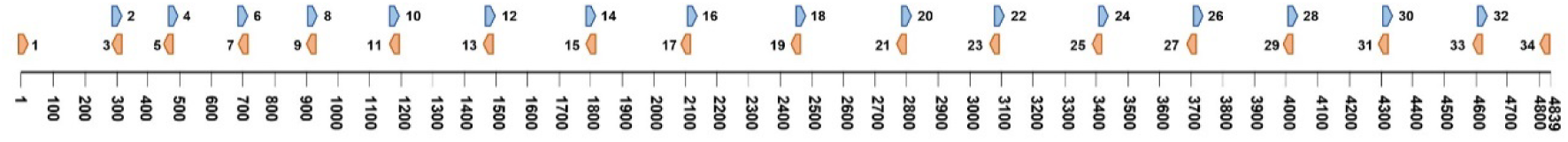
Schematic representation of the primer pairs used for *GmCad1* sequencing. The numbers below the line represent the nucleotide positions. The numbers next to each primer representation refer to Supplementary Table 1

**Figure S2.**
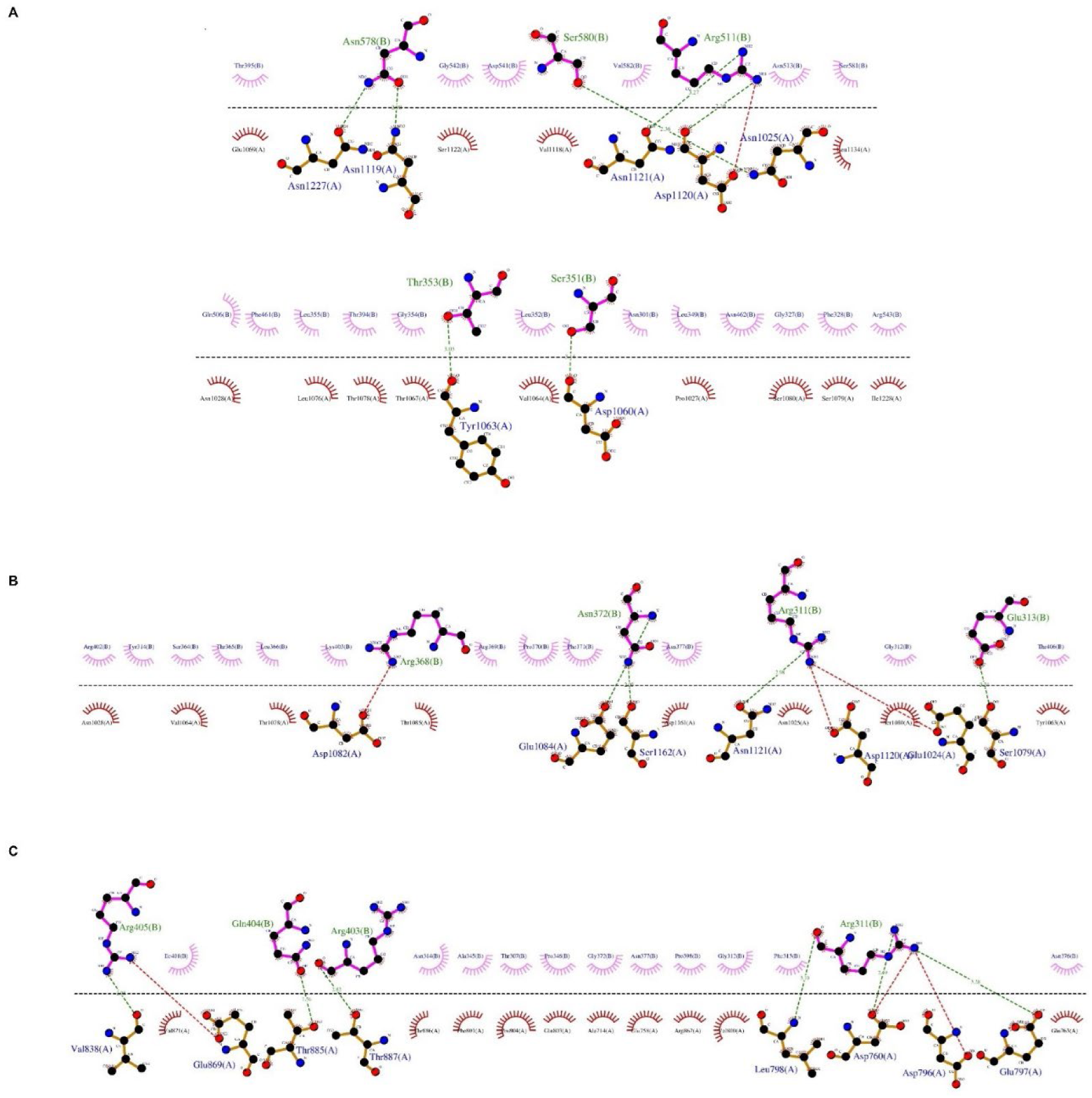
LigPlot+ interaction diagram of G. molesta CR7-11 and Cry1A proteins at 5 pH. (A) Cry1Aa (B) Cry1Ab (C) Cry1Ac. In green are shown the residues of Cry1A proteins and in blue) the residues of CR7-11 that are involved in the interactions. Green dashed lines show the hydrogen bonds and their distance, and the red dashed lines show the hydrophobic interactions.

**Figure S3.**
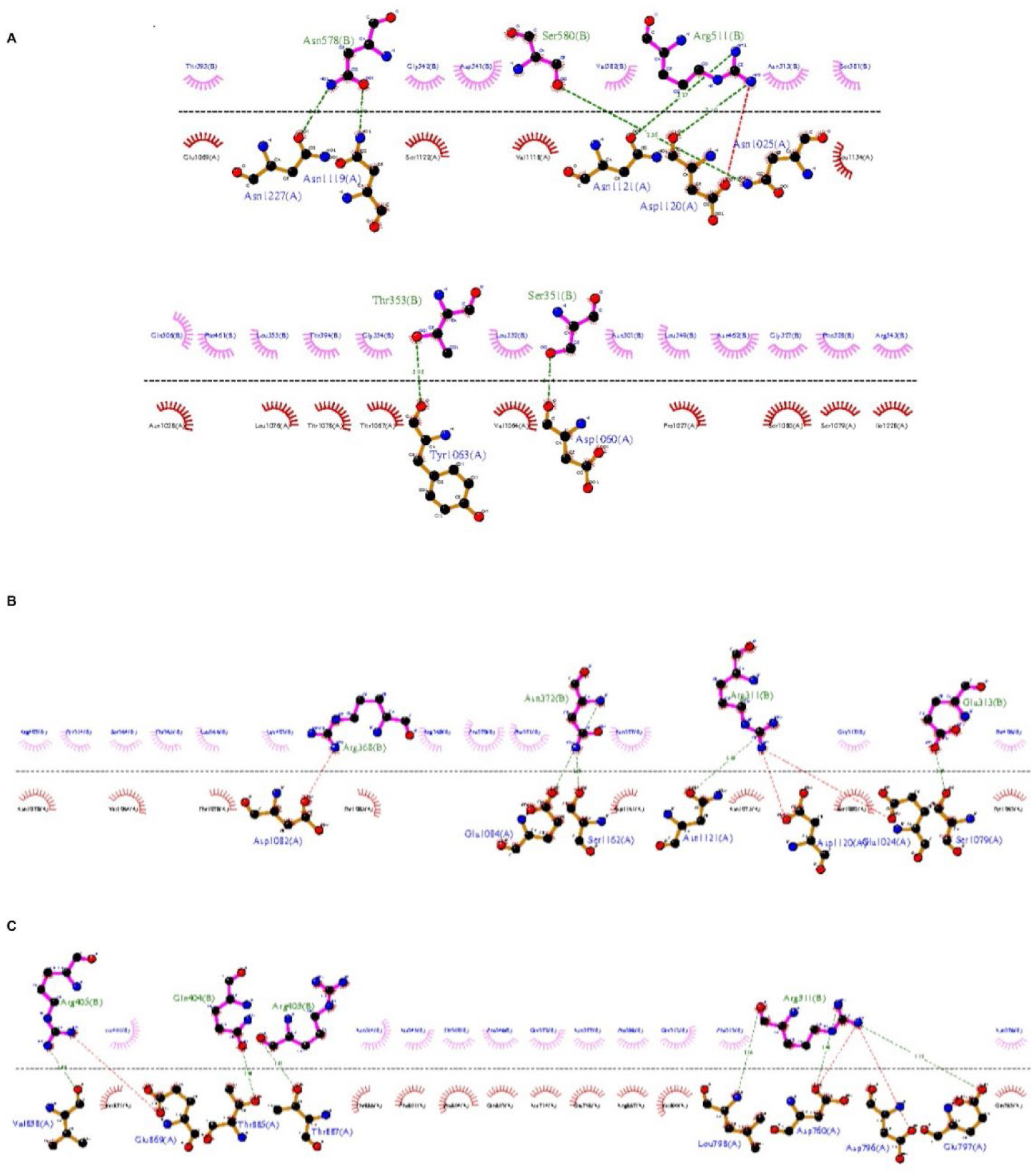
LigPlot+ interaction diagram of G. molesta CR7-11 and Cry1A proteins at 7.5 pH. (A) Cry1Aa (B) Cry1Ab (C) Cry1Ac. In green are shown the residues of Cry1A proteins and in blue) the residues of CR7-11 that are involved in the interactions. Green dashed lines show the hydrogen bonds and their distance, and the red dashed lines show the hydrophobic interactions.

**Figure S4.**
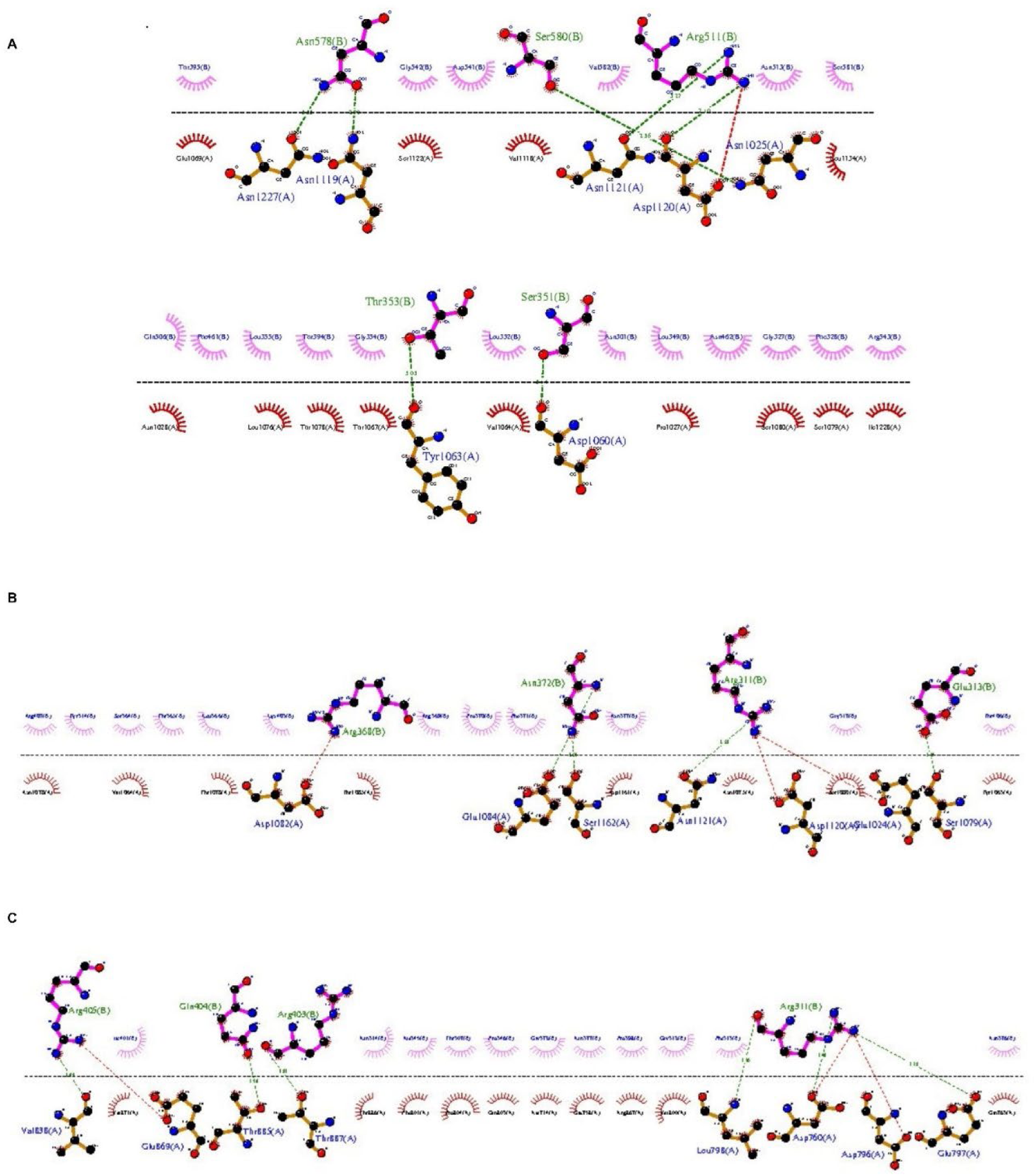
LigPlot+ interaction diagram of G. molesta CR7-11 and Cry1A proteins at 10 pH. (A) Cry1Aa (B) Cry1Ab (C) Cry1Ac. In green are shown the residues of Cry1A proteins and in blue) the residues of CR7-11 that are involved in the interactions. Green dashed lines show the hydrogen bonds and their distance, and the red dashed lines show the hydrophobic interactions.

---

**Supplementary Table 1.**
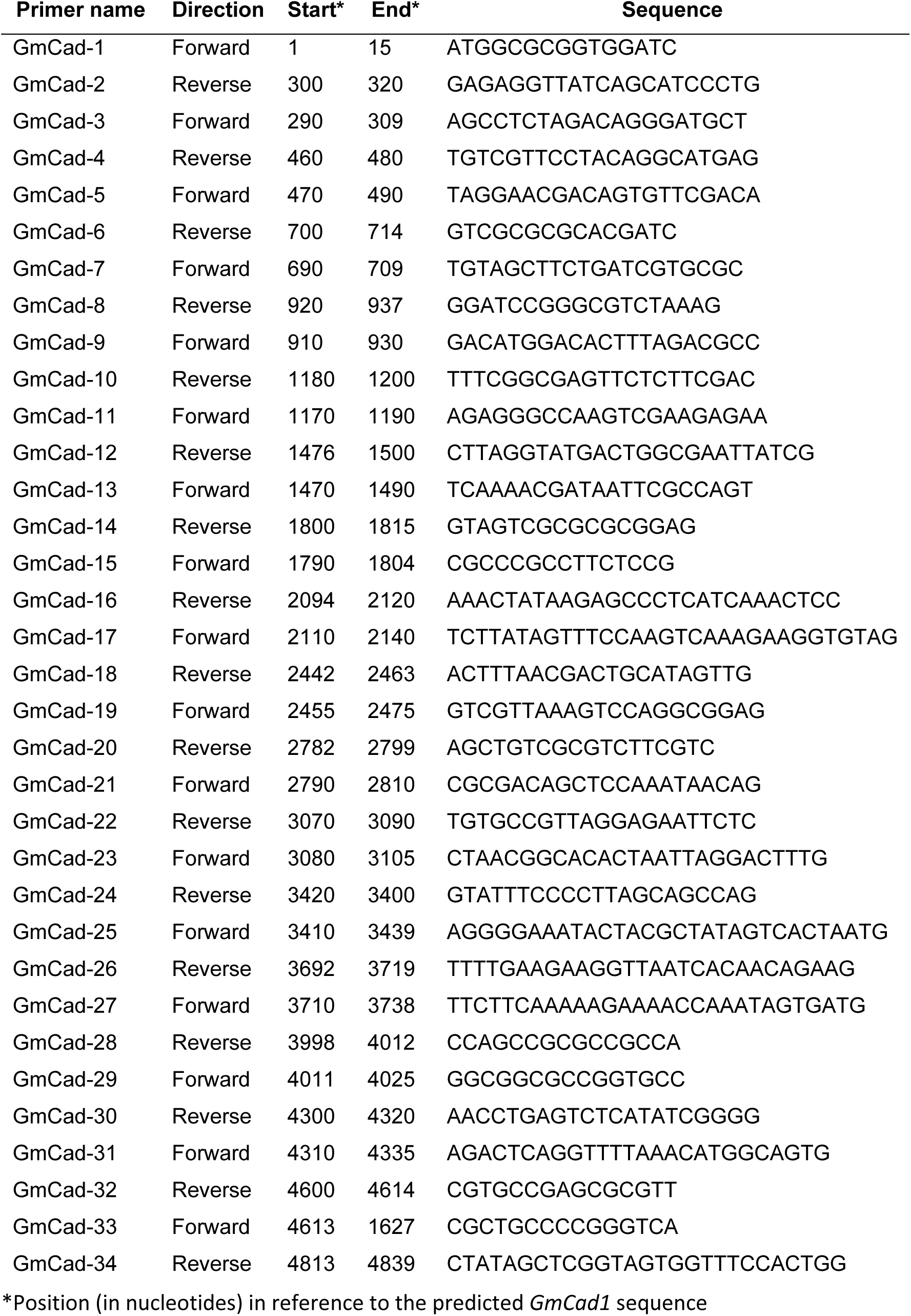
Primer pairs used for *GmCad1* amplification and sequencing.

